# GetOrganelle: a fast and versatile toolkit for accurate *de novo* assembly of organelle genomes

**DOI:** 10.1101/256479

**Authors:** Jian-Jun Jin, Wen-Bin Yu, Jun-Bo Yang, Yu Song, Claude W. dePamphilis, Ting-Shuang Yi, De-Zhu Li

**Affiliations:** Germplasm Bank of Wild Species, Kunming Institute of Botany, Chinese Academy of Sciences, Kunming, Yunnan 650201, China; Center for Integrative Conservation, Xishuangbanna Tropical Botanical Garden, Mengla, Yunnan 666303, China; Kunming College of Life Sciences, University of Chinese Academy of Sciences, Kunming, Yunnan 650201, China; Center of Conservation Biology, Core Botanical Gardens, Chinese Academy of Sciences, Mengla, Yunnan 666303, China; Department of Biology, The Pennsylvania State University, University Park, PA 16801, United States; Southeast Asia Biodiversity Research Institute, Chinese Academy of Science, Yezin, Nay Pyi Taw 05282, Myanmar

**Keywords:** Assembly, Assembly graph, Plastome, Organelle genome

## Abstract

GetOrganelle is a state-of-the-art toolkit to assemble accurate organelle genomes from NGS data. This toolkit recruit organelle-associated reads using a modified “baiting and iterative mapping” approach, conducts *de novo* assembly, filters and disentangles assembly graph, and produces all possible configurations of circular organelle genomes. For 50 published samples, we reassembled the circular plastome in 47 samples using GetOrganelle, but only in 12 samples using NOVOPlasty. In comparison with published/NOVOPlasty plastomes, we demonstrated that GetOrganelle assemblies are more accurate. Moreover, we assembled complete mitogenomes of fungi and animals using GetOrganelle. GetOrganelle is freely released under a GPL-3 license (https://github.com/Kinggerm/GetOrganelle).

## Background

Organelles contain plastid (including chloroplast and other forms of plastid) and mitochondrial genomes, also called as plastome and mitogenome. Most plastomes, around 120-150 kb in size, maintain a conserved circular and quadripartite structure, with a pair of inverted repeat regions (IRs) that separate a large single copy (LSC) and a small single copy (SSC) regions [1, 2]. Mitogenome exists in all eukaryotic organisms and shows high diversity in genome size and forms. To date, six main types of mitogenome organization have been recognized [3]. Of them, animals have single circular mitogenome from 11-28 kb in size (type I); and fungi and plants have single circular mitogenome with introns from 19-1000 kb in size (type II), or a circular large molecule from 20-1000 kb in size and small circular plasmid-like molecules (type III), or homogenous linear molecules from 1-200 kb in size (type V). So far, organelle DNA markers have been the most widely used for phylogenies [4-8] and DNA barcoding [9-12]. Since the rapid advances of high throughput sequencing technologies, sequencing cost has decreased tremendously in recent years. Due to high number of copies of organelle genomes in a single cell, it is feasible to get enough reads to assemble complete organelle genomes from the low coverage whole genome sequencing (WGS) data [13, 14].

To date, in spite of untested assembly quality, there are 2937 plastomes and 9217 mitogenomes (including 8128 animals, 563 fungi, 60 plants, and 255 protists) available in GenBank (https://www.ncbi.nlm.nih.gov/genome/browse#!/organelles/, accessed on April 15, 2019). Many processes or pipelines assembling organelle genomes have been described. For example, SPAdes [15], SOAPdenovo2 [16], and CLC Genomics Workbench (https://www.qiagenbioinformatics.com/) have been widely used to assemble the WGS data, after which the organelle genomic scaffolds/contigs were selected or filtered out using a reference genome for the further concatenation [17] or post assembly gap filling and closing [18, 19]. However, those approaches are computationally intensive with limited probability of organelle genome completeness [20]. The IOGA (Iterative Organellar Genome Assembly) pipeline [21] incorporated Bowtie2 [22], SOAPdenovo2, SPAdes and other dependencies for recruiting plastid-associated reads and conducting de novo assembly. But, the plastomic scaffolds/contigs need to be finalized by other programs. The ORG.asm [23] and NOVOPlasty [24] were reported as fast tools to conduct de novo assembly of a complete organelle genome. However, both tools had not been systematically compared until a recent preprint work was published [25]. Freudenthal *et al.* [25] presented a benchmark of seven chloroplast assembly tools and found significant differences among those assemblers. In their test, our toolkit GetOrganelle (https://github.com/Kinggerm/GetOrganelle) significantly outperformed all other assemblers in consistency, accuracy and successful rate. Nevertheless, Freudenthal *et al.* [25] only tested a limited application scope and limited parameter range of GetOrganelle. Besides, an organelle genome such as a plastome usually has flip-flop configurations or other isomers mediated by repeats [26-28], which was not covered by all above mentioned tools except GetOrganelle, nor by this benchmark test [25].

GetOrganelle toolkit supplies numbers of scripts and libraries for recruiting target organelle reads from the WGS data, manipulating and disentangling assembly graphs, and generating reliable organelle genomes accompanied with labeled assembly graph for user-friendly manual completion and correction (Figure 1). The from-reads-to-organelle process could be finished using the “get_organelle_from_reads.py” script with a single line command, which serves as the main workflow of GetOrganelle. This script exploits Bowtie2, BLAST [29], SPAdes, as well as Python library numpy, scipy, and sympy as dependencies. It starts with recruitment of initial target-associated reads by using Bowtie2 and taking target genome(s) or fragment(s) sequence(s) as the seed; the initial target-associated reads (seed reads) will be treated as “baits” to get more target-associated reads through multiple extension iterations, which is similar to those of the MITObim [30] and IOGA [21] pipelines. However, the core algorithm of this script for reads extension is using the hashing approach, which cuts the reads into substrings with certain length, called “words”, and adds them to a hash table, called “accepted words” (or “baits pool”). During each extension iteration, the “accepted words” is dynamically increasing as new target-associated reads being cut and added as “words”. Then, the total target-associated reads are *de novo* assembled into a FASTA assembly Graph (“FASTG”) file using SPAdes. Non-target contigs in the FASTG assembly are further automatically identified and trimmed by their connections, coverages and BLAST hit information using a target-gene-based database. The slimmed FASTG file is used to calculate all possible paths of the complete target organelle genome based on the graph characteristics and the coverages of the contigs. In some cases, when the assemblies could not be solved as any kind of circular path or was too complicated circular path(s) for GetOrganelle to solve, GetOrganelle will export the target contigs/scaffolds. Meanwhile, the slimmed FASTG file can also be visualized by Bandage [31] to manually finalize the complete target genome or export the assembled target contigs/scaffolds. In this study, we illustrated the mechanism and workflow of GetOrganelle, tested GetOrganelle with a wide range of both parameters and samples, compared the assemblies of NOVOPlasty, the assemblies of GetOrganelle, and the published plastomes in details.

**Figure 1.**
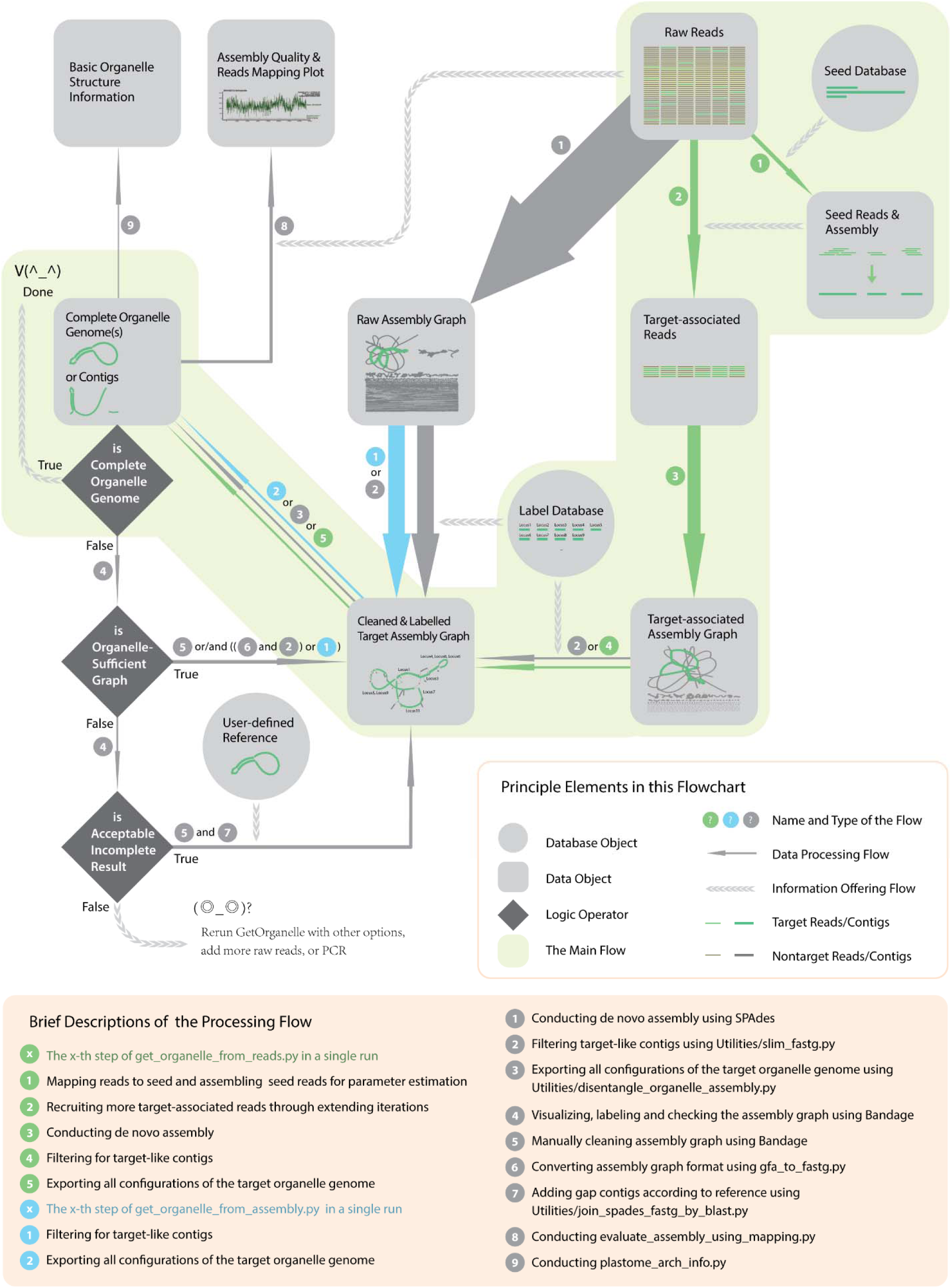
The workflow of GetOrganelle toolkit. The thumbnails inside data objects show an example of plastome assembly. The solid arrows denote the data processing flows and associated directions, with their width proportional to general computational burden. All green solid arrows together describe a complete run from reads to organelle genome(s), which can be finished using get_organelle_from_reads.py with one-line command.

## Results

### Performance of GetOrganelle

In the test of performance, we assembled reads of an angiosperm species, *Haberlea rhodopensis* Friz. [GenBank Sequence Reads Archive accession number (SRA): SRR4428742], using the complete plastome or a short plastomic fragment *rbcL* gene of a gymnosperm species, *Gnetum parvifolium* (Warb.) W.C. Cheng [GenBank accession (GBK): NC_011942.1] as the seed (Figure 2, Additional file 1: Figure S1, Additional file 2: Table S1). The runs with all tested word size ratios (defined as word size value over effective mean read length), except for 0.9, assembled an identical complete plastome, no matter the initial seed was a complete plastome or a short plastomic fragment *rbcL* gene, no matter the pre-grouping was enabled or not.

**Figure 2.**
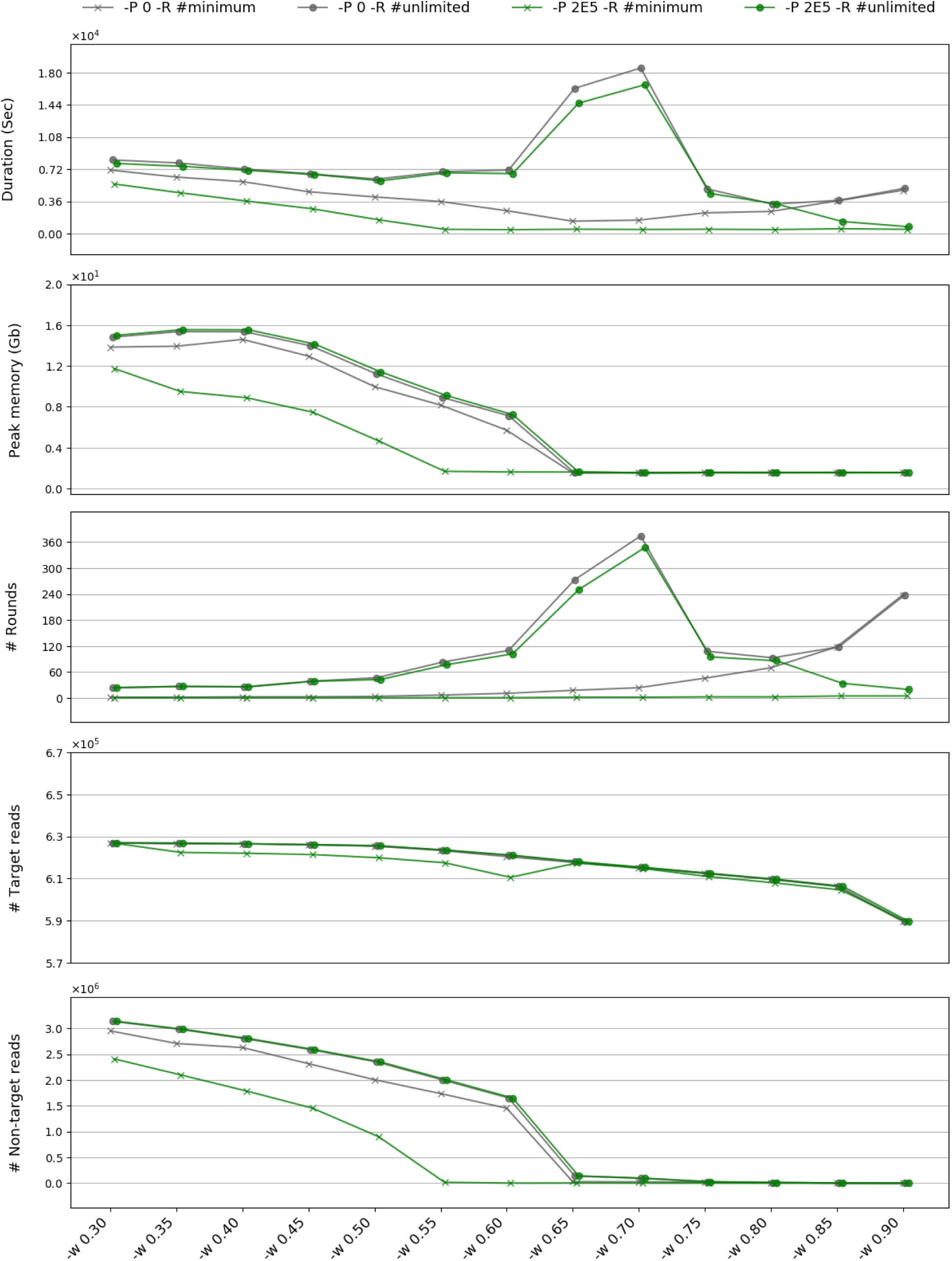
Assessing the plastome assembly performance characteristics of GetOrganelle using online reads of an angiosperm species, *Haberlea rhodopensis* (SRA: SRR4428742) as the dataset, using the complete plastome of a gymnosperm species, *Gnetum parvifolium* (GBK: NC_011942.1) as the seed. The assessment was conducted across a range of word size ratios (after the flag “-w” as labelled at the bottom). The green marks denote using the pre-grouping value 200,000 (after the flag “-P”), while the grey marks denote that the pre-grouping was disabled. The cross marks denote using the minimum rounds of extension iterations for achieving a complete plastome or stabilizing the incomplete plastome result, while the solid circle marks denote using unlimited number of rounds (“-R 1000” in practise). From the top to the bottom, of assembling raw reads into the same complete plastome (or incomplete in all runs of “-w 0.90”), those five subgraph present the total computational time cost by seconds, the maximum memory cost by gigabytes, the actual number of rounds GetOrganelle took, the number of target plastid reads GetOrganelle recruited, the number of non-target reads GetOrganelle recruited.

As the word size ratio (defined as word size value over the effective mean read length) set from 0.3 to 0.9, the running duration of GetOrganelle varies in different conditions (Figure 2), while the maximum memory occupation generally decreased when word size ratio was smaller than 0.65 and remained unchanged when it reached 0.65 (Figure 2). A too large word size, such as 0.9 in our test, would risk an incomplete result. The time cost of the ones with unlimited rounds (grey marks in Figure 2) generally showed a W-shape curve. The time cost of the ones with minimum rounds and pre-grouping disabled (green cross marks in Figure 2) generally showed a shallow-V-shape curve. The time cost of the ones with minimum rounds and pre-grouping (green triangle marks in Figure 2) generally showed a L-shape curve.

When testing with unlimited number of rounds, the ones with pre-grouping enabled (grey triangle marks) and the ones with pre-grouping disabled (grey cross marks) recruited exactly the same number of the final accepted reads (Figure 2) thus the same final assemblies, and similar computational times and memory costs. A few exceptions that the pre-grouping ones cost less time happened when the word size ratio value is high (Figure 2). However, considering the minimum rounds situation, the pre-grouping ones generally needed significantly much less rounds (<6) of extension for recruiting most target-associated reads for downstream achieving a complete plastome or stabilizing the incomplete result (Figure 2). Extending after five rounds is spent mostly in non-target reads. By cutting off the latter unnecessary rounds in the minimum rounds (green marks), the pre-grouping ones (green triangle marks) cost significantly less memory when word size ratio < 0.65 and always cost less time, even significantly sometimes (Figure 2).

### Reassembling plastomes from 50 published datasets using GetOrganelle

For assembling 50 plant datasets (Additional file 2: Table S2), as the word size ratio increased from 0.6 to 0.8, both the mean memory and the mean duration for assembling the 50 samples reduced, and the number of circularized genomes decreased as well (Figure 3). For runs with auto mode, which used an automatically estimated word size for reads extension, the computational usage of both mean memory and the mean duration ranked the most intense, while the number of obtained circularized genomes are 39 that ranked the best in the tested parameter sets (Figure 3). Eight more samples obtained the complete circular plastome by adjusting the word size for extension or *k-*mer for SPAdes, or other parameters (see details at https://www.github.com/Kinggerm/GetOrganelleComparison). GetOrganelle exported all potential configurations mediated by potential flip-flop recombination induced by IR or other shorter repeats.

**Figure 3.**
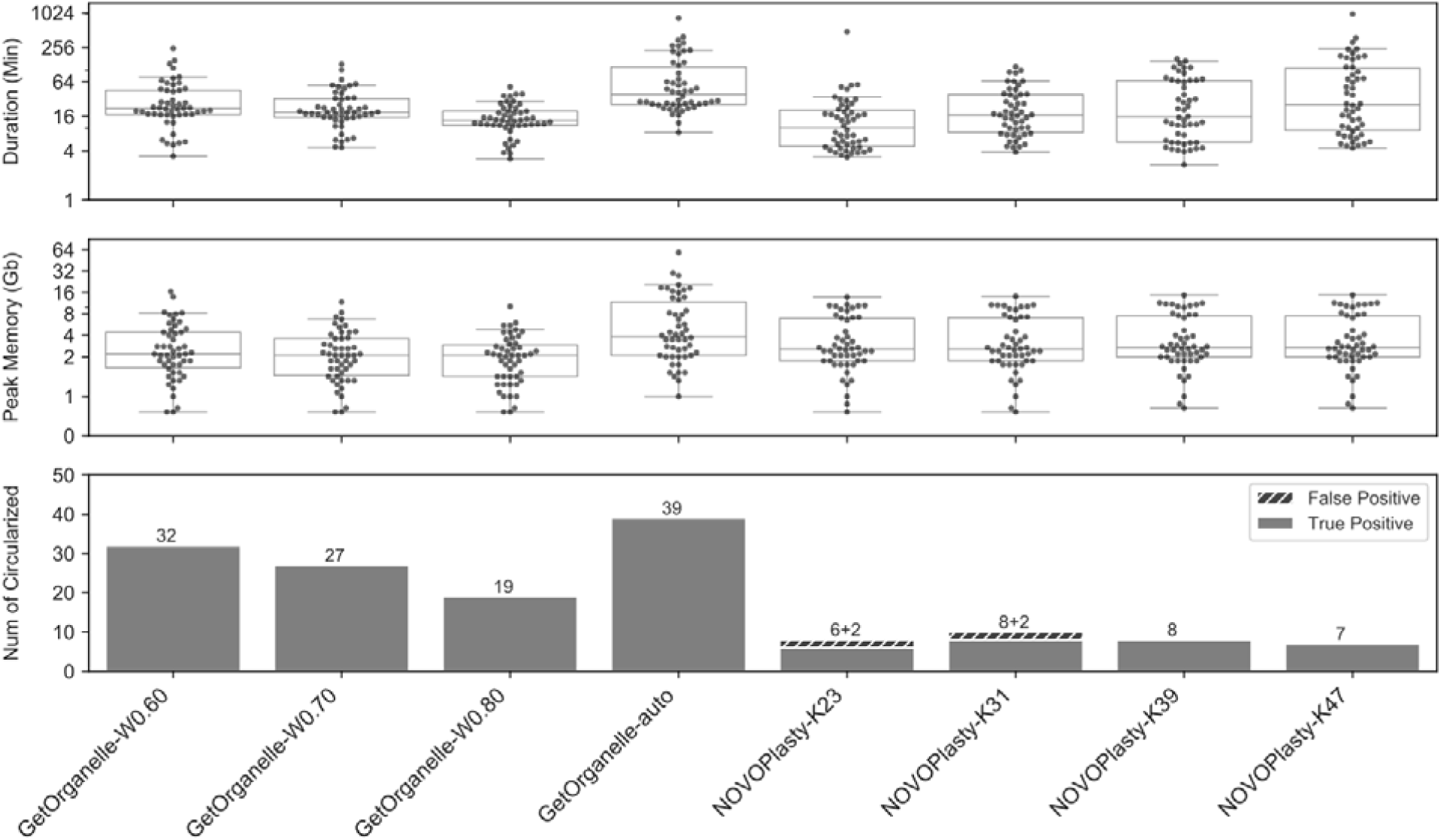
Comparisons of four sets of runs using GetOrganelle and four sets of runs using NOVOPlasty on assembling 50 public datasets. The labels in the bottom denote the program and key arguments for each set of 50 runs. The dots in the upper boxplot presents time cost by seconds of finishing each run. The dots in the middle boxplot presents the maximum memory usage by gigabytes of finishing each run. The bottom histogram presents the number of circularized genomes for each set.

**Figure 4.**
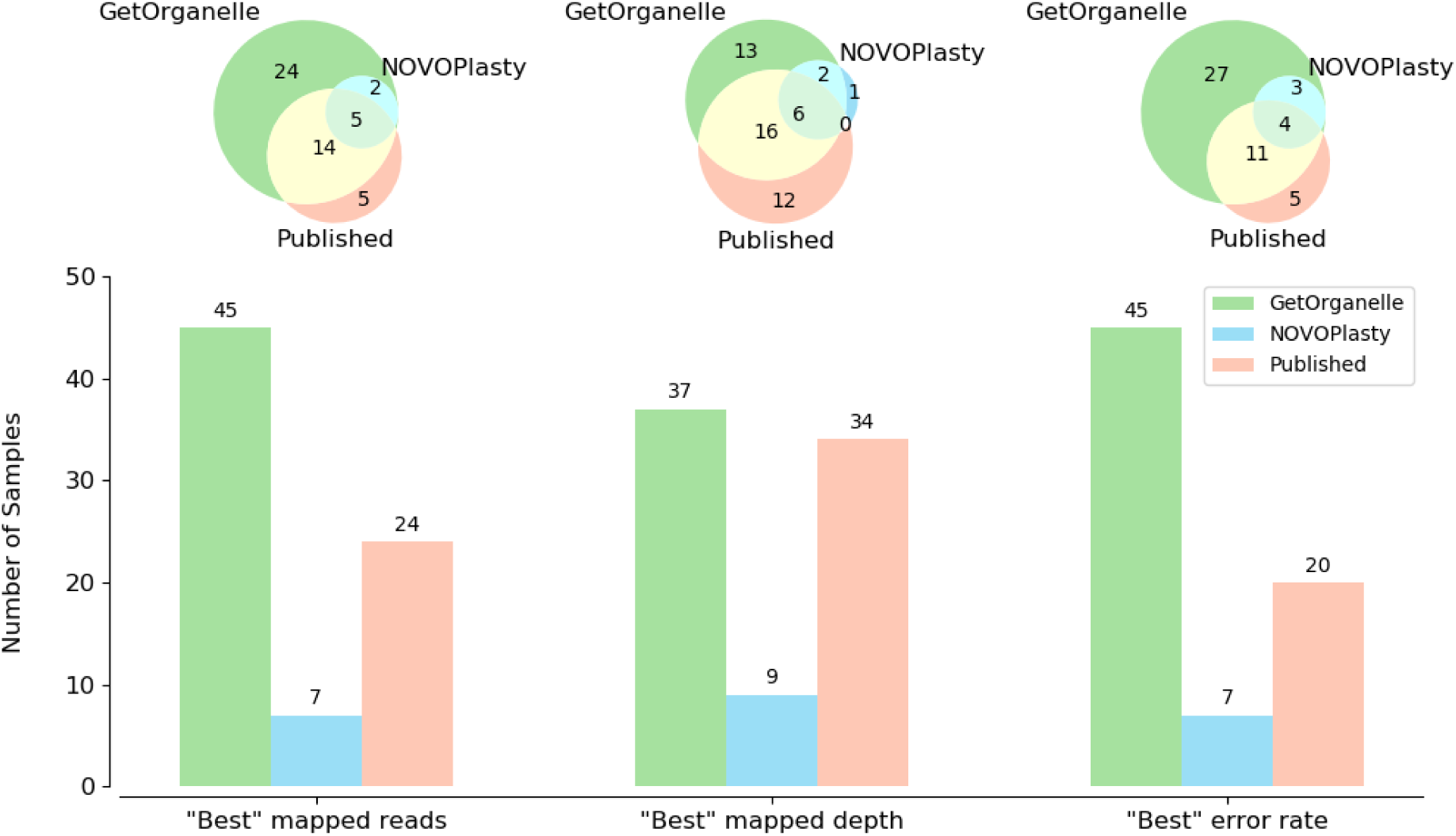
Evaluating assembly qualities of GetOrganelle plastomes, NOVOPlasty plastomes, and published plastomes using reads mapping.

When the complete circular plastome was obtained, GetOrganelle generally outputted the consistent result based the same raw reads with different parameters. In the plastome test, there are 33 samples that obtained the complete circular plastome reassembled from multiple runs using different parameters of GetOrganelle. Twenty-five out of 33 samples have identical results based on different parameters. Five out of 33 samples (SRA: SRR5602586, SRR5602590, SRR5602594, SRR5602605, SRR6932851; we use the SRA number to represent the sample thereafter) have results that diverged in 1 or 2 bp indels. For SRR5602577, the plastome of the GetOrganelle-W0.8 run is shorter than the customized run in a 6-bp poly-A/T indel. For SRR5028199, the plastome of the GetOrganelle-W0.6 run is longer than the plastome of any other 4 runs in a 19-bp repeat (“GAAAAGAAAGAATGAGAAA”). For SRR5602597, the plastome of the GetOrganelle-W0.8 run is shorter than the plastome of any other 5 runs in a 36-bp repeat and a 273-bp indel/repeats.

Finally, GetOrganelle reassembled the complete circular plastome(s) in 47 of the 50 samples. In the remaining three species, two species (*Ginkgo biloba* L., ERR2206741; *Salvinia cucullata* Roxb., SRR6478596) had two break points in the LSC/SSC region, and one species (*Musa balbisiana* Colla, SRR2057084) consisted of plastome fragments (see details at https://www.github.com/Kinggerm/GetOrganelleComparison).

In 14 samples, the GetOrganelle-reassembled plastomes contained identical sequence to the published ones (Additional file 2: Table S2, Table S3). In 26 samples of the rest, the minimum difference of GetOrganelle and published ones were fewer than 5 site-differences and/or less than 100 bp-differences (Additional file 2: Table S2). Most of differences are repeats or indels. Reads mapping values generally supported the GetOrganelle plastome rather than the published ones (Additional file 2: Table S3). For example, the IR boundary regions of *Laurus nobilis* L. (SRA: SRR5602602; GBK: KY085912.1) which showed significant assembly differences between the published one and GetOrganelle-reassembled one, had smooth mapping plot for GetOrganelle plastome but drastically fluctuated mapping plot for the published one (Additional file 1: Figure S2). Those differences can be also monitored from the summarized mapping quality (Additional file 2: Table S3), where the GetOrganelle plastome had similar matched depth (221.60 vs 221.65) with a much smaller deviation (31.43 vs 37.88), as well as the smaller error rate (1.16% vs 1.17%) with a much smaller deviation (1.11% vs 1.56%).

### Reassembling plastomes from 50 published datasets using NOVOPlasty

For assembling the same 50 datasets (Additional file 2: Table S2), the average duration of NOVOPlasty increased as the *k-*mer values increased from 23, through 31 and 39, to 47, while the average memory cost was very close among the four *k-*mer values (Figure 3). The duration of NOVOPlasty was generally shorter than GetOrganelle when using *k-*mer values 23 and 31, and was similar to GetOrganelle-W0.8 when using the *k-* mer value 39, and was only shorter than GetOrganelle-auto when using the *k-*mer value 47. The mean memory cost of the NOVOPlasty runs was lower than GetOrganelle with word size ratio 0.60, 0.65, and auto.

Concerning the completeness rate of NOVOPlasty, only 15 samples were “claimed circular” (i.e., eight for *k-*mer 23, ten for *k-*mer 31, eight for *k-*mer 39, and seven for *k-*mer 47) (Figure 3, Additional file 2: Table S3). Of the claimed circular samples, three samples have gaps or ambitious sites in K23 runs, six samples have gaps or ambitious sites in K31 runs, five samples have gaps or ambiguous sites in K39 runs, and five samples have gaps or ambitious sites in K47 runs. Also, the claimed circular samples might not be really circularized, such as that there are 2 contigs in the K31 run of SRR5602581 and in the K47 run of SRR5602590. Some claimed circularized samples (such as the K23/K31 runs of SRR2037123, the K23 run of SRR5602581) were problematic, because the assembled lengths are highly deviated from the published ones and reassembled ones using GetOrganelle. In the remaining 12 samples that NOVOPlasty obtained the reasonable lengths, the mapping quality of NOVOPlasty assemblies ranked well, although generally not better than those of GetOrganelle especially in the error rate (Additional file 2: Table S3).

Besides, the assemblies of NOVOPlasty are largely inconsistent and varied severely when using different *k-*mer values. Using NOVOPlasty, there are nine samples that obtained the complete circular plastome reassembled from multiple different runs. However, only three samples (SRA: ERR1917165, SRR5602589, and SRR5602602) out of the nine obtained identical plastomes with published.

### Assembling mitogenomes using GetOrganelle

For fungus mitogenome test, GetOrganelle successfully reassembled the complete circularized mitogenome in 24 out of 50 datasets (Additional file 2: Table S4). The average duration for those tests is 10,428 s, ranging from 843 s (SRA: SRR5803928, circularized) to 41,311 s (SRA: SRR5804145, circularized). The average memory cost for those tests is 12.1 G, range from 2.9 G (SRA: SRR5804137, circularized) to 40.1 G (SRA: SRR5804145, circularized). The sizes of circularized results range from 33,655 bp (SRA: SRR5803909), to 209,881 bp (SRA: SRR5804018).

For animal mitogenome test, GetOrganelle successfully reassembled the complete circularized mitogenome in 29 out of 56 datasets (Additional file 2: Table S5). Those successful samples include three samples that were excluded by MitoZ for “mitochondrial derived reads ratios of 0”, although our tests used 1∼5 folds bases of the tests in MitoZ paper [14]. Of the 56 datasets, GetOrganelle failed in calling any animal mitogenome contigs in five samples. The average duration for those tests is 10,428 s, ranging from 843 s (SRA: SRR5803928, circularized) to 41,311 s (SRA: SRR5804145, circularized). The average memory cost for those tests is 12.1 G, range from 2.9 G (SRA: SRR5804137, circularized) to 40.1 G (SRA: SRR5804145, circularized).

## Discussion

### Seed and label database in GetOrganelle

For assembling the plastome from the reads of an angiosperm *Haberlea rhodopensis* with high and homogeneous-distributed plastid coverage, using the plastome of a gymnosperm species *Gnetum parvifolium* (Warb.) W.C.Cheng as the initial seed is feasible. For the same set of reads, even using a short-conserved DNA fragment (*rbcL)* as the seed that is sufficient to recruit almost all target-associated reads to assemble a complete plastome, but less efficient. The reason is that the seed is only used for reads mapping to collect the initial reads (“baits” or seed reads), after which the seed will no longer be used (Figure 1). GetOrganelle then use the seed reads to recruit the target-associated reads by extension with multiple rounds, which is mainly based on the nature of the read overlaps with the seed reads (Additional file 1: Figure S1), but not a reference-based assembly process. Nevertheless, using the organelle genome of a non-related species as the seed can only be applied to plastome seed recruitment, but not animals nor fungi mitogenome seed recruitment dues to their highly divergent evolution rate (See more in the “mitogenomes of fungi and animals” part).

Since the label database (see Figure 1) in GetOrganelle is used to provide supervision signal for labeling contigs before semi-supervised learning, the label database is preferred to be conserved regions such as coding regions and not necessary to be the intact set of genes. After labeling, GetOrganelle would use an integrated strategy as described in methods to approach identifying target contigs, which largely replies on the contig coverages and assembly graph characteristics. Thus, the label database as gene fragments has limited influence on the final organelle-complete graph decision and would barely affect the scaffolding of organelle contigs.

### Word size in GetOrganelle

The value of word size, like the *k-*mer in De Bruijn graph-based assembly, is important to the feasibility and efficiency of read extension. But it differs from *k-*mer in its unique role of overlap threshold for data filtering. The smaller the word size is, the lower the threshold would be. The use of word size does not only examine the nature of connection between any two target reads, but also in a way judge the connection strength by the coverage. Even if the plastome does share homologous region with mitogenome or nuclear genome, the reads of those non-target genomes would not be largely recruited due to the low coverage, meaning that any two reads of those non-target reads are less likely to be connected in the shallow coverage WGS, or genome skimming data.

The word size is generally negatively correlated to the memory usage (Figure 2), which is opposite to the situation of *k-*mer. The memory cost decreased because of the larger the overlap threshold causing the less reads connected and recruited based on the overlap, and thus smaller word space in extension and smaller *k-* mer space in *de novo* assembly. Thus, the *k-*mer only influences the memory with the same data, and the word size influences both memory and the numbers of reads (i.e. the data size) to be recruited. For many real datasets, because the plastome might share homologous regions with the non-target genomes, a small word size meaning a relaxed threshold would not promptly stop the extension when the extension was walking through the non-target genomes. In the case of Figure 2, it showed wasting the computational resources to use a word size ratio less than 0.65.

The word size affects the duration of a whole assembly run from different aspects (Figure 2). Generally, without the pre-grouping, a larger word size has the faster speed of extension across the candidate reads that reduced the time cost. While a small word size will recruit more non-target reads, by contrary, the time cost will be increased. Besides those two antagonistic effects, the calculation times for read comparison reduces with increasing the word size. Without round limitations and without the pre-grouping, the time cost showed a complicated “W-shape” curve with increasing the word size (grey marks). When it comes to using the minimum rounds of extension without the pre-grouping, there would be a word size value that costs the least time (green cross marks). In the pre-grouping and minimum rounds situation, the time cost is nearly constant in the common-used range, but it remarkably increased with reducing the word size ratio from 0.65 to 0.3 (green triangle marks). In empirical studies, although the minimum number of rounds is not known in advance, with the pre-grouping algorithm, most extension process within 5 rounds (a very few cases with extremely low plastome coverage requires about 15) recruited the target reads covering the whole plastome, which has been specified as the minimum rounds of extension.

The best word size varies with read length, read quality, total number of reads, percentage of organelle DNA content, heterogeneity of organelle base coverage, and other factors. If there is no user-assigned word size value, “get_organelle_from_reads.py” will automatically estimate an optimal word size value using a set of empirical-customized functions based on the data characteristics. The automatically estimated word size tends to be a small value to enhance the successful rate, which may increase the computational burden (Figure 3). Meanwhile, the automatically estimated word size does not always guarantee the best performance (see eight plastome samples with customized parameters in our tests).

### *K*-mer in GetOrganelle

One advantage of GetOrganelle uses SPAdes as the de novo assembler, which combines the assemblies from multiple *k-*mers. In our 50-sample test, the most successful assemblies are the largest *k-*mer that the data permit, which is 127 for read length ≥150 bp or 91 for read length up to 100 bp. This would be curtly and speciously owing to that the base coverage is usually high enough to use the largest *k-*mer as possible for assembling the complete plastome or mitogneme from the genome skimming data. However, except for those with low base coverage for plastome (e.g. SRR5602610, ca. 14×), the assembly of plastome still suffered from the base coverage heterogeneity issue or the read error, with which there would be some regions that large *k-*mer cannot overlap. For example, if only one large *k-*mer value (i.e. 91 for read length ca. 100 bp and 127 for read length ≥ 150 bp) was used for each run, some samples (tests not shown) would not assemble a complete circular plastome; By using the *k-*mer gradient, those samples achieved a complete plastome at the same large *k-*mer.

Another advantage of GetOrganelle to use a multi-*k*-mer supported assembler is that GetOrganelle could iteratively try to disentangle the assembly graph of each *k-*mer from the largest to the smallest, then find a larger one with the organelle-sufficient graph (See the definition in the Method). A larger *k-*mer value will be preferred when there are longer repeats and coverage is sufficiently high. However, when the largest *k-*mer used in the analysis could not help achieving the complete plastome, the assembly graph of a smaller *k-*mer would be checked automatically. The “GetOrganelle-auto” runs of SRR5602587 and SRR5028199 benefit from this design.

### Computational consumption of GetOrganelle

Freudenthal *et al.* [25] showed that GetOrganelle has moderate efficient in both time and memory usage among tested assemblers. However, they only used the default options that is inclined to have high organelle genome completion rate rather than high computational efficiency. For studies that does not aim at achieving complete organelle genomes, researchers could easily adjust the options such as increasing the word size or turning on “--fast” to notably speed up and reduce memory usage, and still keep both the successful rate and assembly quality at a higher level than those of other assemblers. However, we do not recommend users to do so. Firstly, for current version of GetOrganelle, a circularized result could better ensure the assembly quality than the incomplete result. Secondly, extremely high word size also generates higher error rate with two cases in our 50 samples, such as the GetOrganelle-W0.8 runs of SRR5602577 and SRR5602597.

### Accuracy of the published/reassembled plastomes

Concerning the chromosomal structure level assembly accuracy, the GetOrganelle-reassembled plastomes contained identical sequence to the published ones in 14 samples, including *Selaginella kraussiana* (Kunze) A. Braun (SRR2037123) that was reported to have large directed repeats (DRs) [32]. Reads mapping plot of GetOrganelle reassembled complete or near complete plastomes are generally smooth with little unusual peaks. In contrast, the NOVOPlasty reassembled ones (i.e. ERR964904, SRR2037123, SRR5602581 as mentioned above) had unreasonable shorter lengths, but claimed them to be circularized. According to the reads mapping plot (https://www.github.com/Kinggerm/GetOrganelleComparison/tree/master/eval/Published), many published ones (such as SRR1145775, SRR2057084, SRR5602601, SRR5602602, SRR6478596, SRR7630500) have misidentified IR boundaries or contig overlaps as revealed by the extremely unusual peaks.

Concerning the depth of the assembly, according to the evaluation using reads mapping (Additional file 2: Table S3), 37 assemblies using GetOrganelle have the best-ranked matched depths (smaller average and smaller deviation). The rest 13 assemblies (3 of them were not circular) have also values very close to the best-ranked ones except for one sample (SRR2037123) for which NOVOPlasty produced a problematic result with aberrantly high coverage, and two samples (SRR2057084, SRR6478596) for which GetOrganelle did not generate a circular plastome.

Concerning the nucleotide level assembly accuracy, according to the evaluation using reads mapping (Additional file 2: Table S3), 45 assemblies using GetOrganelle have the best-ranked error rate (smaller average and smaller deviation), the rest 5 assemblies (even 3 of them were not circular) have also values very close to the best-ranked ones. However, many published plastomes (e.g. SRR5602575-SRR5602578, SRR5602581-SRR5602584, SRR5602587, SRR5602588, SRR5602592, SRR5602593, SRR5602595, SRR5602597, SRR5602598, SRR5602599, SRR5602600, SRR5602609, SRR5602610, SRR6932851, SRR7630500) have nucleotide sites that incongruent with the consensus of most reads as revealed by extremely high mismatch/indel points.

Our result also showed that GetOrganelle generally outputted the consistent result using the same raw reads with different parameters. As mentioned above, 31 out of 33 samples have identical or almost identical (<= 2 bp difference) results based on different parameters. There are mainly three reasons for the discrepancies in the rest few samples. Firstly, a large word size would recruit significantly less reads even cause incomplete result (such as -w 0.8 in SRR5602587 and -w 0.9 in the performance test). Secondly, there are small repeats that limited reads cannot precisely identify, which is another reason for -w 0.8 in SRR5602587. Thirdly, there are polymorphism in the data, such as SRR5602587 recovered by the auto mode.

Our result also suggested that using reads mapping qualities is an easy and important way for evaluating organelle genome assemblies. Besides, we tested the 50 samples here not only for showing that GetOrganelle has better accuracies, but also appeal to all plastome providers for the necessity of making raw data available to the public (e.g. SRA), for making reproducible, evaluable and amendable assemblies, which will be benefit to future comparative analyses of organelle genomes [33].

### Large Inverted, Directed Repeats and “weird” plastomes

The entanglement of repeats is one of the challenges in organelle genome assembly, which was not addressed in reported plastome assemblers. The largest repeat pair in a canonical plastome can be the IR. Given that flip-flop recombination mediated by IR is regular [34, 35], a genuine *de novo* assembler such as GetOrganelle should export at least two configurations when IRs exist. Even if we assume that there was only one configuration in vivo, short sequencing reads theoretically cannot tell between those two configurations when IRs exist. However, NOVOPlasty produces, if successfully, only one representative of the plastome structure at the risk of misleading unconversant users into referring SSC orientations as an important inversion.

Another issue concerning IR is the identification of the IR boundaries. A traditional assembly method which use the CLC or any other assembler to make the contigs and finish the circular based on a reference is prone to create a plastome with a similar IR length/boundary to the reference. That was why PCR verification were needed for the four boundaries of a canonical quadripartite plastome after “assembly”. This obsolete concept still shadowed lots of empirical-study scholars in the new age. However, a graph-based method for finishing the plastome circle utilizes the original contig connections supported by the reads. Provided sufficient coverage (e.g. 100×), there would be no need for PCR verification nor reads-mapping. A rhetorical question for some recalcitrant opponents is why not use PCR to verify any other parts of the plastome since they are also assembled by short reads. The straightforward answer to the opponents comes from the accurate configuration result of GetOrganelle. GetOrganelle correctly recovered the extreme contraction or loss of a large pair of IRs in *Juniperus cedrus* Webb & Berthel. (SRR1145775) and two *Picea* species (ERR268390, SRR5028199). Another interesting example is that GetOrganelle successfully assembled the plastome of *Selaginella kraussiana* (SRR2037123) with two DRs when the assembler was not given the prior knowledge of IR nor DR. The reassembled sequence is identical to the published one (MH549643.1), which is a verified assembly using PCR [32].

It is a challenge to assemble some “weird” plastomes (e.g., plastome reduction, gene translocations, and IR expansion, or contraction, or loss) using some traditional methodologies [20]. For example, non-photosynthetic plants have reduced plastomes and many gene translocations [20, 36, 37]. If *de novo* assembly cannot yield a complete sequence of the plastome, gap filling and PCR verification are needed to close the gap of the contigs/scaffolds [20, 38]. In our tested species, we successfully assembled a complete plastome for an extremely reduced plastome in a holoparasite *Cytinus hypocistis* (L.) L. (ERR964904) using a customized strategy. Meanwhile, many holoparasites from Balanophoraceae, *Cuscuta* spp. (Convolvulaceae), Lennoaceae, and Orobanchaceae, as well as hundreds of hemiparasites were assembled circular plastomes (W.-B. Yu et al., unpublished data). For the IR lacking species, like some Fabaceae species, a single circle of the assembly can be displayed by Bandage, which provides the evidence for exempting the PCR verifications.

### Mitogenomes of fungi and animals

In our tests, assembling animal mitogenomes, generally cost much more computational resources, in terms of both duration and memory than assembling plastomes and fungi mitogenomes, even the animal mitogenomes are much smaller. One reason for the increase of computational burden could be the increase of data size, which is the number of bases. Another reason may be the inaccurate target organelle coverage estimation, therefore improper word size estimation. Most of the time, GetOrganelle underestimated the target organelle coverage, generated extremely small word size values, and extended too much non-target reads. The underestimation of target organelle coverage may be caused by the limitation of the default seed database (see more discussion below). It is also worth mentioning the relatively low animal mitogenome percentage (ca. 0.5% in average) than plastome percentage (ca. 3% in average) and fungi mitogenomes (ca. 3% in average) in the tested genome skimming data, although the underlying relationship between the target organelle percentage and the increased computational cost was untested.

The mitogenomes of fungi and animals usually have much higher nucleotide substitution rate [39, 40]. Therefore, unlike assembling plastomes, a relatively closely related seed and label database would be indispensable for assembling and identifying mitogenomes using the extension and assembly strategy of GetOrganelle. If GetOrganelle failed with default database for animals or fungi mitogenome assembly, it is suggested that users rerun the same sample with their own database. The limitation of the default seed database or the default label database may be responsible for some failures. For example, SRR1946581 has 1,796,709 reads (ca. 225 Mbp) mapped to the default seed database, but none of its resulting contigs hit the label database, because the default label database is currently a small near-random subset of the default seed database.

### Other small repeats, broken graph and manual completion

Aiming at solving the organelle genome structure, other small repeats caused the awkward tangles in the assembly graph. Plastomes of some plant clades were reported to have multiple short repeats, such as some clades in Ericaceae [38, 41], Geraniaceae [42], and Pinaceae [27, 43].

One common category is the tandem repeat, which can be visualized as whirls in the assembly graph. T The main problem for disentangling the tandem repeats in an assembly graph is detecting the multiplicity (copy number) of the contigs. While many bacterial genomes assemblers used the minimum or the greedy algorithm for detecting multiplicity, which is simply assuming every tandem repeat contig has multiplicity two and constructing the assembly as a path that traverses the repeat twice [44-46]. GetOrganelle disentangles the tandem repeats based on both the connection information and the contig coverage, which has broader realistic sense if the target organelle genome has some tandem repeats of more than 2 copies and the reads offers high coverage to overcome the error (Additional file 1: Figure S3). Other repeats such as the inverted repeats other than the large IR, do not only cause the assembly graph tangles, but also might mediate the recombination and produce isomers in the same individual [26-28, 47]. For researchers studying plastome rearrangement and recombination dynamics, GetOrganelle serves as the right tool to exhaust the limits of the assembly graph and present all possible configurations, such as plastomes of *Juniperus cedrus* (SRA: SRR1145775) and *Picea* species (ERR268390, SRR5028199), mitogenomes of fungi species (SRR5801935, SRR5804018) and animals species (SRR4340274, SRR136494). Currently, those potential configurations mediated by short repeats are suggested to be further confirmed by read pair information of mapping or PCR verification. As the development of long insert size library or long-read sequencing (e.g. PacBio), these data can be added for repeat resolution and configuration confirmation. A function that could use the long library reads or long-read sequencing data and estimate the proportion of all the candidate configurations is expected in future versions of GetOrganelle.

There are still some assembly graphs that are not sufficient (broken graph) or too complicated to estimate the multiplicity. In that case, users could conduct the manual completion or manual multiplicity with the simplified assembly graph using Bandage. The concomitant cognominal TAB-formatted file could supply as important information source for manual completion or manual multiplicity. Another advantage of manual completion is feasible to detect interchange among nuclear genome, mitogenome and plastome. Especially when multiple organelle genome mode was chosen for plant data, a tangly assembly graph mixed with plastome, mitogenome, and mitogenome-plastome-shared contigs would be achieved.

## Conclusions

GetOrganelle is a fast and versatile toolkit for de novo assembly of complete and accurate organelle genomes using WGS data. According to our tests, we have demonstrated that the GetOrganelle toolkit can efficiently and accurately assemble different types of organelle genomes from broad range organic lineages. In general, comparing with reassembled results from NOVOPlasty, GetOrganelle has far better successful rate for assembling plastomes while consuming close computational resources. Noteworthily, GetOrganelle reassembled plastomes generally have much higher accuracy than reassembled ones by NOVOPlasty and published ones that assembled by multiple tools in according with our reads mapping evaluation. Besides, GetOrganelle could generate all possible configurations when a plastome or mitogenome has flip-flop configurations or other isomers mediated by repeats.

Potential applications of GetOrganelle include quickly extracting organelle genomes from whole genome assemblies and evaluating organelle genome qualities. Assembling organelle genomes from metagenomic and transcriptomic data would be also possible given a customized database and scheme. The maximum extending length option enables rough control of length of target assembly, which could be used to quickly assemble interested loci or genes from the metagenomic and transcriptomic data. Additionally, the Python Classes and Functions defined in GetOrganelleLib could be used to manipulate and disentangle non-organelle assembly graphs.

Currently GetOrganelle exports all possible configurations without library information of the paired-end reads. However, the long insert size library or long-read sequencing data can be used for repeat resolution and configuration confirmation. A function that could use this information and estimate the proportion of all the candidate configurations is expected in future versions of GetOrganelle. Improvements in default database construction are also expected, for better parameter estimation and mitogenome successful rate.

## Methods

### Workflow of organelle genome assembly using GetOrganelle

GetOrganelle v1.6.2 consists of two major scripts (“get_organelle_from_reads.py” and “get_organelle_from_assembly.py”) and 17 minor scripts (under the directory “Utilities”; for processing or evaluating organelle assemblies) (Figure 1), a series of libraries (under the directory “GetOrganelleLib”; including default seed database, default label database, and Python Classes/Functions), and dependencies (under the directory “GetOrganelleDep”; including Bowtie2, SPAdes, BLAST+). The major script “get_organelle_from_reads.py” could pipe all 5 steps described below (also see green solid arrows in Figure 1) together using a single line command to assemble organelle genome(s) from raw reads. The major script “get_organelle_from_assembly.py” could extract organelle genome(s) from assembly graphs (Step 4-5; also see blue solid arrows in Figure 1) that can be assembled using SPAdes [15] or Velvet [48].

### Step 1. Mapping reads to seed and assembling seed reads for parameter estimation

The initial step of assembling target organelle genome(s) from reads via GetOrganelle (“get_organelle_from_reads.py”) is using Bowtie2 to map reads to seed sequence(s), which can be complete reference organelle genome(s) or organelle fragment(s) (Figure 1, Green solid arrow 1). Currently, the default seed of GetOrganelle (under the directory “GetOrganelleLib/SeedDatabase”) covers embryophyte plastomes, non-embryophyte plastome, embryophyte mitogenomes, embryophyte nuclear ribosomal DNA, animal mitogenomes, and fungal mitogenomes (in the option referred as “embplant_pt”, “other_pt”, “embplant_mt”, “embplant_nr”, “animal_mt”, “fungus_mt” separately). The generated mapped reads is called seed reads here (stored at *.initial.fq).

The seed reads will be treated as “baits” to recruit more target-associated reads in the next step. In the “auto” mode (see the last paragraph of Step 2 below), the seed reads will be also coarsely assembled into seed contigs, which will be used for parameter estimation in Step 2.

### Step 2. Recruiting more target-associated reads through extending iterations

After creating “baits”, GetOrganelle (“get_organelle_from_reads.py”) recruits new target-associated reads by comparing candidate reads to “baits” and updating the “baits” with overlapped new reads (Figure 1, Green solid arrow 2). In this extension process, the key comparison method for determining overlaps is the classic substring hashing. Substring(s) is specified as “word(s)” here to discriminate the “*k*-mer”, another term with similar concept in the assembly process. The uniform length of the “words” is thus named as “word size”.

Before the core extension iterations, “get_organelle_from_reads.py” will create an index and assign unique ids for each set of duplicated reads to avoid repeatedly calculating (information stored at file “temp.indices.1”). Those reads with duplicates will be also utilized for downstream “Pre-grouping”. “Pre-grouping” is an algorithm for speeding up target reads recruitment. This algorithm is based on the idea that it would be more efficient to firstly compare reads that are more likely to be target-associated. Given that the organelle genomes usually have more copies hence higher base coverage than most non-organelle chromosomes, the duplicated reads are more likely to be organelle-associated than non-duplicated reads. “get_organelle_from_reads.py” will group a certain number of duplicated reads (after option “-P”) into groups based on read overlap information using the same substring hashing method mentioned above. Any group, including those with only a single read, will have a hash table storing words chopped from all reads of this group. Any two groups sharing at least a single word in their hash table will be merged. After “Pre-grouping”, a group resembles a set of connected pseudo-contigs (information stored at file “temp.indices.2”). During following extension iterations, once a read was accepted as a target-associated read, all other reads (ids) in the same group will be marked as acceptable.

The “get_organelle_from_reads.py” begins the core extension iterations with constructing a hash table, by cutting the initial reads (“baits”) into “words” and adding those initial “words” to the hash table, named as “Accepted Words” (AW). If a “word” of a candidate read hits the AW, the index of this read will be marked as accepted (Accepted Index, AI), with all words of this read being immediately added to AW. As mentioned above, all other read ids in the same group will be treated as accepted. In this way, AW increases contingent on meeting an acceptable read which overlaps previously accepted reads or be in the same group with previously accepted reads. During each iteration (round), “get_organelle_from_reads.py” goes through all candidate reads one by one to check whether a read is acceptable. After a user-specified number of rounds or when no new read been accepted in a complete round, “get_organelle_from_reads.py” will stop the extension process and output all accepted reads (stored at file “filtered_*.fq”). For low memory machines or testing purpose, accepted reads per round can be outputted separately (followed with flag “--out-per-round”) along with AW dumped after each round.

The value of the word size (followed with flag “-w”), like the *k-*mer in assembly, is crucial to the feasibility and efficiency of this process. The best word size changes upon data and will be affected by read length, read quality, total number of reads, percentage of organelle reads, heterogeneity of organelle base coverage, and other factors. In the “auto” mode when there is no user-assigned word size value, “get_organelle_from_reads.py” will automatically estimate a proper word size with a set of empirical-customized functions, based on the data characteristics.

### Step 3. Conducting *de novo* assembly

The recruited target-associated reads will be then automatically assembled using SPAdes (Figure 1, Green solid arrow 3). Both paired and unpaired reads will be used. The outputs of each *k-*mer of SPAdes include an assembly graph (FASTG format), which records the connections of contigs as a graph with some allelic polymorphism and assembly uncertainty. Other assemblers generating the assembly graph, such as Velvet, is also feasible for completing this step but is not implemented in GetOrganelle yet. The intermediate results are stored in the subfolder “filtered_spades”.

### Step 4. Filtering for target-like contigs

Due to the frequently occurred sequence similarities in some regions or sharing copies among plastome, mitogenome, and nuclear genome, the accepted reads from Step 2 usually unavoidably contain lots of non-target reads (Figure 1, Green solid arrow 4 and blue solid arrow 1). As a consequence, the output assembly graph might also contain some non-target contigs. This is one of the problems that are not well addressed nor solved by all previously reported tools.

In this step, GetOrganelle uses BLAST against a local label database (see next paragraph), contig connection information and contig coverages to search for the target-like contigs from the original assembly graph file. By conservatively deleting non-target contigs, GetOrganelle would output a simplified assembly graph file, along with a concomitant cognominal TAB-formatted file (file name uses the postfix “.csv” to be in conformity with Bandage) recording the hitting details. This step is automatically fulfilled by the two major scripts or can be separately executed using the script “slim_fastg.py”.

For GetOrganelle, the local label database of a certain organelle is made from the coding regions of that organelle genome. The default local label database covers the same six organelle/region types as the seed database. A contig that hit the target organelle database will be labeled with gene details (information stored at the TAB-formatted) and called target-hit-contig here. Any contig that be in the same connecting component with a target-hit-contig, which means directly or indirectly connecting to that target-hit-contig, is called a target-associated-contig. Specifically, under the embplant_pt or embplant_mt mode, GetOrganelle will by default keep both plastome and mitogenome contigs for downstream clustering contigs by coverage. The processing of slim_fastg.py is designed to be conserved to avoid removing true target contigs of plant plastome or mitogenome.

### Step 5. Exporting all configurations of the target organelle genome

In this step, GetOrganelle uses the simplified assembly graph file and the BLAST hitting table produced by slim_fastg.py to: a) further accurately identify target organelle contigs, b) estimate multiplicities (copy number) of contigs, and c) export all possible distinctive path(s) [stored as FASTA file(s)] from the organelle assembly graph (stored as a cognominal GFA format file) (Figure 1, Green solid arrow 5 and blue solid arrow 2). Each possible path is a possible configuration of the target organelle genome. This step is fulfilled by the two major scripts or can be separately executed using the script “disentangle_organelle_assembly.py”. In case of organelle genome with a large number of repeats in a few samples, to avoid generating inexhaustible combinations, GetOrganelle sets up an option for limiting the calculating time of disentangling. For the major scripts, when it failed to export circular sequence(s) from the assembly graph in the ways mentioned below or ran out of time limit, it would try a second run to export the contigs, which would be mainly the target-hit-contigs.

There are three definitions needs to be clarified:

**Definition 1:** an organelle-sufficient graph is an assembly graph with contigs completely covers one complete organelle genome.

**Definition 2:** an organelle-only graph is an assembly graph only with true contigs of one organelle genome.

**Definition 3:** an organelle-equivalent graph is an organelle-sufficient graph and an organelle-only graph.

There are three main assumptions for GetOrganelle to disentangle the assembly and declare the result as complete circular organelle:

**Assumption 1:** All configurations, if there are more than two, of the target organelle genome are compositionally identical. This assumption limits the multiplicities of contigs to be the same among different configurations. In the other words, polymers found in real plastid DNA molecules [49], whereas GetOrganelle only exported the monomer form, and potential sub-genomic configurations are currently not implemented in the current version. If there are parallel contigs caused by nucleotide polymorphism, all subgraphs composed of any of those polymorphisms will be disentangled independently. Therefore, all configurations of each subgraph will be compositionally identical.

**Assumption 2:** The topology of each configuration of each organelle genome is a single circular molecule. This assumption holds when the real organelle genome is a circular molecule or organized in polymers (most plastomes, and type I and type II mitogenomes) and the assembly graph is an organelle-sufficient graph. If this assumption is violated, GetOrganelle only export the target contigs.

**Assumption 3:** The coverage values of contigs of the same organelle genome are generally proportional to their multiplicity (copy number). The coverage values of contigs with the same multiplicity of the same organelle genome generally held consistent.

#### a) Identifying target organelle contigs

Only using the BLAST hit information to identify target organelle contigs is risky. For example, some contigs, including mitochondrial contigs that have short length of similar region to plastid genes and target-like shallow-depth contigs from contamination, would be target-hit-contigs but not true target contigs (false positive). Some are true target contigs but too short or diverged from sequences in label database to be target-hit-contigs (false negative). Therefore, we used additional information to improve the identification of target contigs, such as the assembly graph characters (the assumption 1 and 2) and contig coverage values (the assumption 3). GetOrganelle uses an integrated strategy that iteratively uses all or part of following modules to approach this task.

1. Using the BLAST hit information to roughly cluster contigs with target labels. In detail, GetOrganelle firstly calculates a customized hit weight value (HW) for each BLAST hit record. For those records hitting the same gene in the local BLAST database, only the record with the best HW will be kept as the only valid hitting record for that gene. The HW of a hit record is simply defined as the product of the hitting length of the query (HL) and query contig coverage (QD) (HW = HL * QD). Given our experience that the false positive hits generally correspond to shorter length and shallower depth contigs, this definition could largely exclude the false positive hit records. Each gene in the BLAST database would thus be aligned to no more than one contig in the assembly graph. Secondly, GetOrganelle calculates a customized contig weight value (CW) of a certain organelle type for each contig. For example, the plastid CW for a contig is defined as the summation of the HWs of all plastid gene hit records of that contig, while the mitochondrial CW for the same contig is defined as the summation of the HWs of all mitochondrial gene hit records of the same contig. For a contig, if the target CW is much (default factor: 3 times) larger than the non-target organelle CW, this contig would be labeled as a target-anchor contig (very likely to be a true target contig), vice versa. If the difference is smaller than the factor (default factor: 3 times), the contig would have multiple labels. Using the HW and CW, GetOrganelle roughly eliminates most of “noisy” and non-target contigs, even with the false positive hits.
2. Adding more target labels to some target contigs that do not hit the database according to assembly graph characters. Based on the assumption 2, any configuration of the target organelle genome is a single circular molecule. As a result, in an organelle-sufficient graph, both the two ends of any true target contig should be connected to at least one true target contig. Based on this deduction of the assumption 2, if the tail end of a true target contig, Contig A (marked as A_tail_) has only one edge that connects A_tail_ and the head end of another unknown contig, Contig B (B_head_), then Contig B should be a true target contig. However, in a real assembly graph with missing contigs (incomplete organelle genome), Contig B may be missing and the unknown contig connected to A_tail_ may be a non-target contig. In considering of the complicated situation in a real assembly graph, only when Contig B is in-between two target-anchor contigs (Contig A and Contig C) with sequence (A_tail_-B_head_-B_tail_-C_head_), and A_tail_ only connect to B_head_ and C_head_ only connect to B_tail_, GetOrganelle labels Contig B as a target-anchor contig with CW=0.
3. Using coverage values of contigs to remove contigs with coverage value that significantly deviates from the target-anchor contigs. Based on the assumption 3, GetOrganelle uses the Gaussian mixture distribution to approximate the coverage values of all contigs in the simplified assembly graph, which is a mixture of different organelle contigs and nuclear contigs. In most cases of empirical plant genome skimming data, the plastome has significantly higher coverage than mitogenome, the coverage of which in turn is higher than nuclear genome except highly repeated regions. Therefore, in a plant WGS dataset, the coverage values of plastid, mitochondrial and nuclear contigs are expected to be classified into different components of the Gaussian mixture distribution. So does other mixture situations, such as contig assemblies of a lichen that is a symbiosis with fungi and algae or cyanobacteria. GetOrganelle could thus delete the contigs with coverage value far from the target coverage distribution. Specifically, GetOrganelle applies EM (Expectation-Maximization) algorithm with the semi-supervised learning and the weighted Gaussian mixture model to cluster the coverage values of all candidate contigs. Here the semi-supervised learning means there is labeled data, the coverage values of the target-anchor contigs, which will not be updated during EM iterations. The coverage value of a contig in the Gaussian mixture model is weighted by the length of the contig.
4. Removing contigs isolated from the main target connected component that includes the target-anchor contigs. Based on the assumption 2, true target contigs in an organelle-sufficient assembly graph should be in one connected component. Thus, for a real organelle-sufficient assembly graph, GetOrganelle takes the connected component with most target-anchor contigs and delete other components. In detail, GetOrganelle calculates a customized component target weight value (TW) for each connected component of the assembly graph. The TW of a connected component is defined as the summation of the target CWs of all contigs of that component. Assuming a real graph is an organelle-sufficient assembly graph, GetOrganelle sorts connected components by their TWs, finds the component with the significantly largest TW (which is arbitrarily 100 times larger than the second largest TW by default) and remove the contigs of other components from the assembly graph. If GetOrganelle failed to find a component with the significantly largest TW, disentangling the assembly graph as a circular organelle genome failed and turn on the “linear mode”. In a “linear mode”, when GetOrganelle tries to disentangle the assembly graph as contigs or scaffolds, several components with large TW are allowed to remain. In that case, only components with TWs 10000 times (by default) smaller than the largest TW will be removed.
5. Removing tip contigs. A tip contig is a contig with one (in the lateral case) or both (in the linear case) ends that do not connect any other contigs in the assembly graph nor to itself as circular. Based on the assumption 2, an organelle-equivalent graph will not contain any tip contig, because any partial sequence of a circular DNA molecule would have its upstream sequence and downstream sequence. In detail, GetOrganelle will check whether a tip contig is a target-anchor contig before removing it. If the tip contig is a target-anchor contig, it is more likely that the assumption 2 is violated, and in most cases, the assembly graph is not an organelle-sufficient graph or “a broken organelle graph”.

#### b) Estimating multiplicity of contigs in an organelle-only graph

There are two information sources for estimating multiplicities for contigs in GetOrganelle. One is the coverage values of contigs. Any contig in an organelle-only graph would have at least one copy, which form the basic constraints for downstream multiplicity estimation.

Using the python library Sympy, GetOrganelle firstly creates a set of linear equations to characterize the multiplicity relationship between any two connected contigs. In detail, there are mainly four constraints to build this set of equations. First, the multiplicity of a self-loop contig has no constraints based on connections. Second, if Contig A is not a tip contig nor a self-loop contig, the multiplicity of Contig A is equal to sum of the multiplicities of the contigs connected to A_head_, and equal to sum of the multiplicities of the contigs connected to A_tail_. Third, the multiplicity of a tip contig is arbitrarily set to 1 to avoid over-estimating, although which risks failures in solving these equations. Last, in considering of symmetry of large inverted repeats (such as IRs in plastome), the multiplicity of a sequential repeat contig is constrained to be integer times of that of its nearby contigs. This set of equations would be then solved. Multiplicity values of all contigs would be represented as linear expression of several freedom variables. The values of those freedom variables would be further optimized to minimize the summation of the difference between coverage-based multiplicity and the equation-based multiplicity of each contig (Additional file 3: Figure S3).

#### c) Exporting all possible configurations

GetOrganelle then exhaustively search for all possible paths from this organelle-only graph with contig multiplicities. Each configuration combination would be saved as an independent FASTA format file, with the same head name style to manual completion using Bandage [31] (see below). A circular sequence would be marked “(circular)” in the head of the FASTA format sequence. For plastomes with repeats inside the large IRs, there would be 6 paths (Additional file 1: Figure S3; another similar but more complicated example is SRR5602601 with 12 paths). However, in considering of symmetry, only those paths with identical large IRs (path1 & 2 in Figure S3) are more biologically possible paths. In this case, GetOrganelle would mark these results with identical IRs as the first repeat pattern in the file name.

### Step 5’. Manual Completion

In case of failing to automatically export full sequence by detecting insufficient target assembly graph, exhausting time, or exporting too many possible configuration sequences, manual completion is needed to clean “noisy” and non-target contig/scaffold connections (Figure 1, Grey solid arrow 4 and 5). The simplified assembly graph can be visualized using Bandage [31]. Meanwhile, the concomitant annotation “csv” file can be imported into Bandage and added to the graph as labels, which helps to manually identify target-like contigs/scaffolds. The whole sequence could then be manually exported from the cleaned assembly graph using Bandage, or the scripts “slim_fastg.py” and “disentangle_organelle_assembly.py” (Figure 1).

### Assessing the performance characteristics of GetOrganelle

To better present the performance characteristics of GetOrganelle, we applied a combination of a gradient of word size values (as the form of word size ratio, meaning word size value over effective mean read length, i.e. 0.3, 0.35, 0.4, 0.45, 0.5, 0.55, 0.6, 0.65, 0.7, 0.75, 0.8, 0.85, 0.9), unlimited number of rounds and minimum number of rounds, pre-grouping activated (-P 2E5, i.e. using the top 2E5 duplicated reads to conduct the pre-grouping) and disabled (-P 0) to assemble the plastome of a flowering plant *Haberlea rhodopensis* Friv. (Gesneriaceae) using the reduced dataset (500Mb, SRA: SRR4428742), using the complete plastome of a gymnosperm species, *Gnetum parvifolium* (Warb.) W.C.Cheng (GBK: NC_011942.1) as the seed. We additionally use only the *rbcL* gene sequence of *Gnetum parvifolium* (GBK: NC_011942.1) as the seed and the word size ratio of 0.75 to assemble the same dataset. The minimum number of rounds mode is defined as using the minimum rounds of extension iterations for achieving a complete plastome or stabilizing the incomplete plastome result. The minimum number of rounds is beforehand estimated in the following paragraph.

To assess the characteristics of recruited reads per round, “--out-per-round” was chosen to output recruited reads for each round. Then the script “round_statistics.py” was used to assess the increasing cover percent of the organelle genome using a reads mapping approach. Utilizing Bowtie2, this script maps reads of each round to the final assembled plastomes and calculates the percentage of bases in the plastome that are covered by mapped reads over a certain coverage threshold (the defaults were 0 and 10). The minimum number of rounds could then be found around when the percentage of covered bases reaches 100% or stays unchanged and when the final assembly is stabilized. Besides, the script “round_statistics.py” could generate the base coverage across the organelle genome plot for each extension round to visualize the extension process.

### *De novo* assembly of the plastomes of 50 samples using GetOrganelle and NOVOPlasty

To evaluate the working efficiency and assembly success, we selected 50 samples of vascular plants with raw reads achieved in the GenBank database of Sequence Reads Archive (SRA) (Additional file 2: Table S2). The 50 vascular plants representing 42 species of angiosperms (representing eight major clades, 21 orders and 29 families), four species of gymnosperms, three species of ferns, and one species of lycophytes. Noteworthily, raw reads of the 50 samples are associated with published plastome [50-53], which provided a great opportunity to compare the differences between the published plastomes and the newly reassembled plastomes using GetOrganelle. Besides, since 2018, NOVOPlasty has received more than 100 citations for assembly chloroplast genome in Google Scholar (accessed 1 February 2019) and became one of the most widely used tools for plastome assembly. We thus reassembled 50 samples using NOVOPlasty for comparisons.

The data resources are paired-end reads. The read length varied from 100 bp to 300 bp (Additional file 2: Table S2). In all tests, if the tested data was less than 10,000,000 reads for each end, we used all the reads; if the data was more than 10,000,000 reads of each end, we only select the first 10,000,000 reads for each end. A large raw data may consume more time for recruiting the target-associated reads, and de novo assembly using SPAdes.

We set up four testing groups, i.e., three groups with different word size values (w = 0.6, 0.7, 0.8) (i.e. GetOrganelle-W0.6, GetOrganelle-W0.7, GetOrganelle-W0.8), and an auto-estimated word size group (i.e. GetOrganelle-auto). The extending rounds of all tests were set to 10. All other options are set to default, including the seed. Because incomplete assemblies are unsuitable for comparing mapping qualities in the next part, we additionally added extra runs for eight samples, in which GetOrganelle-auto could not achieve complete plastomes, with customized options (GetOrganelle-customized) for mapping quality comparison. The detail information of commands is presented in additional file 1. The memory usage and time cost of all the tests were recorded.

The same reads of the 50 samples were also assembled by NOVOPlasty using four *k-*mer values, i.e., 23, 31, 39 and 47. The config file of NOVOPlasty was downloaded from the GitHub (https://github.com/ndierckx/NOVOPlasty/blob/master/config.txt), with “Type” as “chloro”, “Genome Range” as 15000-180000, “Save assembled reads” as “yes”, “Seed Input” as the same seed as running GetOrganelle, “Read Length” as the mean read length of each sample (seed Additional file 2), and other parameters unchanged.

### Reads mapping to evaluate plastome assemblies

The script “evaluate_assembly_using_mapping.py” could be used to evaluate circular/non-circular assemblies (Figure 1, Grey solid arrow 8). It uses Bowtie2 to map reads to circular/non-circular assembly, parses the SAM file, counts the number of mapped paired and unpaired reads, counts matched bases for each site (M), mismatched bases for each site (X), insertions between any two sites (I) and deletions for each site (D), and calculates the average and deviation of matched depth, average and deviation of mismatched depth, average and deviation of insertions between any two reference sites, and average and deviation of deletions per reference site for each contig and the whole assembly. If ∑M > 0, a customized error rate would be also calculated as (∑X + ∑I + ∑D)/∑M with deviation var(X + I + D))/∑M. If “--draw” was chose, the script “evaluate_assembly_using_mapping.py” would generate the plot of M/X/I/D at each site/site-interval across the whole assembly using Python library matplotlib. For reproducibility when using the same random seed, a circular assembled sequence would be relinearized to make sure that biologically the same circular plastome would have identical start and end since identical linear sequence. By default Bowtie2 does not support mapping reads to a circular sequence, the script “evaluate_assembly_using_mapping.py” gets around this problem by adding an extra fragment of the head of the original sequence to the tail, and counting the mapping statistics (M/X/I/D) of the sites in the extra fragment back to the statistics of head part of the original sequence.

The script “evaluate_assembly_using_mapping.py” was used to assess all the assembled plastomes. The abnormal characters, such as “*” and “-” in the FASTA-format assemblies of NOVOPlasty were replaced with “N”. NOVOPlasty would also produce multiple completely different sequences with the same sequence name in the assemblies, which would be also modified with different names before assessment. For NOVOPlasty, the evaluation statistics of each sample were based on the best result among those using different *k-*mer values. Here the best result is determined in turn by being true circularized, the largest number of mapped paired reads, largest number of mapped unpaired reads, largest matched depth, smallest error rate, smallest deviation of matched depth, smallest deviation of error rate. For GetOrganelle, except for 8 samples that were generated using customized parameters, other evaluation statistics were based on the assemblies from the “GetOrganelle-auto” runs.

For each sample, when comparing the evaluation statistics of different assemblies, the “best” mapped reads (the hat mark in Table S3) is defined as the largest number of mapped paired reads or equal-largest number of mapped paired reads with largest number of mapped unpaired reads; the “best” mapped depth is defined as the largest matched depth or equal-largest matched depth with smallest deviation of matched depth; the “best” error rate is defined as the smallest error rate or equal-smallest error rate with smallest deviation of error rate.

### *De novo* assembly of the mitogenomes using GetOrganelle

In total, 56 animal samples (the test samples of MitoZ [14], see Additional file 2: Table S4) and 50 fungi samples (Additional file 2: Table S5) were used to test the ability of GetOrganelle to assemble mitogenomes. The mitogenomes of plants have a relative slow nucleotide substitution rate [9], therefore it is feasible for GetOrganelle to achieve sufficient reads to construct the mitogenome-sufficient assembly graph even with a remotely related seed, provided enough coverage. However, there are mainly two challenges for short-read sequencing data to finish the mitogenome architecture. Firstly, there are usually lots of repeats in the plant mitogenome causing the awkward tangles in the assembly graph [54-56]. Secondly, most of plant mitogenomes are not single circular structure, and they can consist of one large circular molecule and small circular plasmid-like molecules (type III), or homogenous linear molecules (type V) [57]. There are frequent horizontal transfers from plastome to mitogenome. In this case, when the coverage of plastome is much higher than that of mitogenome, the multiplicity of shared contigs would be hard to estimate. When the coverage of plastome is similar to that of mitogenome, the parallel contigs in between the shared contigs would be difficult to distinguish. Thus, we did not include plant mitogenome testing due to the general infeasibility nature of assembling complete circularized plant mitogenome from WGS data.

The data resources are paired-end reads. The read length varied from 92 bp to 301 bp (see Additional file 2: Table S4, Table S5). In animal tests, if the tested data was less than 75,000,000 reads for each end, we used all the reads; if the data was more than 75,000,000 reads of each end, we only select the first 75,000,000 reads for each end. The same maximum number of reads for fungi was 15,000,000. All tests were performed with default settings with the *k-*mer values set to 21,43,65,87,127. The detail information of commands is presented in Additional file 3: Text 1. The memory usage and time cost of all the tests were recorded.

## Computer resources for testing

All assemblies were executed on an Intel Xeon CPU machine containing 144 cores of 2.40 GHz, a total of 3TB of RAM, and set up with Linux (Linux version 3.10.0-514.el7.x86 (Red Hat 4.8.5-11)). The testing environment also includes Python v3.6.5, Perl v5.16.3, GetOrganelle v1.6.2, Bowtie2 v2.3.5.1, SPAdes v3.13.0, BLAST 2.2.30+, numpy v1.13.3, scipy v0.19.1, sympy v1.1.1, matplotlib v3.0.2, psutil v5.4.7, and NOVOPlasty 2.7.2 [24]. GetOrganelle is capable of multi-processing in mapping and assembly process, which was disabled in this test for comparable with NOVOPlasty. All tests could be reproduced by accessing https://www.github.org/Kinggerm/GetOrganelleComparison.

## Acknowledgements

We are grateful to Chao-Nan Fu, Han-Tao Qin, Yang Pan, Xiao-Jian Qu, Shuo Wang, Rong Zhang, Fei Zhao and lots of users for giving tests or suggestions.

## Author contributions

J-JJ, W-BY, T-SY, D-ZL conceived the research; J-JJ wrote the script; J-JJ, W-BY analyzed the data; J-JJ, W-BY wrote the draft manuscript; J-JJ, W-BY, J-BY, YS, CWD, T-SY and D-ZL revised and approved the manuscript.

## Ethics approval and consent to participate

Not applicable

